# Burst coding despite unimodal interval distributions

**DOI:** 10.1101/2021.02.18.431851

**Authors:** Ezekiel Williams, Alexandre Payeur, Albert Gidon, Richard Naud

**Affiliations:** Department of Mathematics and Statistics, University of Ottawa, 150 Louis Pasteur, Ottawa, K1N 6N5, Canada; University of Ottawa Brain and Mind Institute, Centre for Neural Dynamics, Department of Cellular and Molecular Medicine, University of Ottawa, 451 Smyth Rd., Ottawa, K1H 8M5, Canada; Institute for Biology, Humboldt-Universität zu Berlin, Berlin, Germany; Department of Physics, University of Ottawa, 150 Louis Pasteur, Ottawa, K1N 6N5, Canada

**Keywords:** neural coding, burst coding, information theory, short-term plasticity

## Abstract

The burst coding hypothesis posits that the occurrence of sudden high-frequency patterns of action potentials constitutes a salient syllable of the neural code. Many neurons, however, do not produce clearly demarcated bursts, an observation invoked to rule out the pervasiveness of this coding scheme across brain areas and cell types. Here we ask how identifiable spike-timing patterns have to be to preserve potent transmission of information. Should we expect that neurons avoid ambiguous patterns that are neither clearly bursts nor isolated spikes? We addressed these questions using information theory and computational simulations. By quantifying how information transmission depends on firing statistics, we found that the information transmitted is not strongly influenced by the presence of clearly demarcated modes in the interspike interval distribution, a feature often used to identify the presence of burst coding. Instead, we found that neurons having unimodal interval distributions were still able to ascribe different meanings to bursts and isolated spikes. In this regime, information transmission depends on properties of the synapses as well as the length and relative frequency of bursts. Furthermore, we found that common metrics used to quantify burstiness were also unable to predict the degree with which bursts could be used to carry information. Our results provide guiding principles for the implementation of coding strategies based on spike-timing patterns, and show that even unimodal firing statistics can be consistent with a bivariate neural code.

## Introduction

The vast majority of neurons in the brain communicate complex and irregular sequences of voltage spikes – a window into neuronal information processing. These spike trains may be parsed into recurring syllables, a small set of short spike timing patterns bearing potentially different meanings. Multiple studies (2–7) have related spike-timing patterns with different types of information. Focusing on the simplest syllables, the burst coding hypothesis separates isolated spikes from isolated bursts of spikes in rapid succession. Alternatively, bursts may merely consist of a number of equally meaningful spikes that neurons sporadically emit at a high-frequency in order to maximize information transmission (8). Determining which coding scheme applies for a given cell type and brain region has important consequences for the interpretation of neuronal responses.

A number of observations support the burst coding hypothesis. First, bursts and non-bursts occur naturally in multiple cell types and brain areas (7, 9–13). Second, bursts both cause (6, 14) and correlate with (7, 15–17) different types of information. Third, a number of cellular mechanisms regulate the generation of isolated bouts of high-frequency spikes (5). Fourth, synaptic mechanisms route bursts and non-bursts dynamically to different targets (18, 19). Lastly, bursts have theoretical advantages for the nervous system as they allow neurons to enhance information transmission (20), distinguish external from internal information (20) and solve the credit assignment problem (21).

A possible pitfall of burst coding is that many neurons do not emit well-separated singlets and bursts, thus potentially restricting burst coding to cells with a clear qualitative demarcation between these patterns. Burst coding has previously been assessed on the basis of the Inter-Spike Interval (ISI) distribution (Fig. 1A) (22–27). A unimodal ISI distribution (Fig. 1Ai) would be expected from a neuron not using burst coding, for instance when spikes are emitted randomly at a specific rate. Alternatively, a bimodal ISI distribution (Fig. 1Aii) suggests that ISIs can be grouped into two distinct modes (Fig. 1Bi-iii): the mode on the left in Figure 1Aii (bottom) corresponding to bursts of short ISIs, and the mode on the right corresponding to relatively well-isolated events. Does a unimodal distribution necessarily exclude a burst code? It is conceivable that neurons may tolerate a blurry separation between bursts and isolated spikes (Fig. 1Ci) if this allows them to communicate more information per spike. For instance, preserving bimodality imposes a strict constraint, which may be detrimental to information transmission (Fig. 1Cii). Thus, we may expect a trade-off between losing information to misclassified bursts/non-bursts and the cost of a restriction on the dynamic range of responses.

**Fig. 1.**
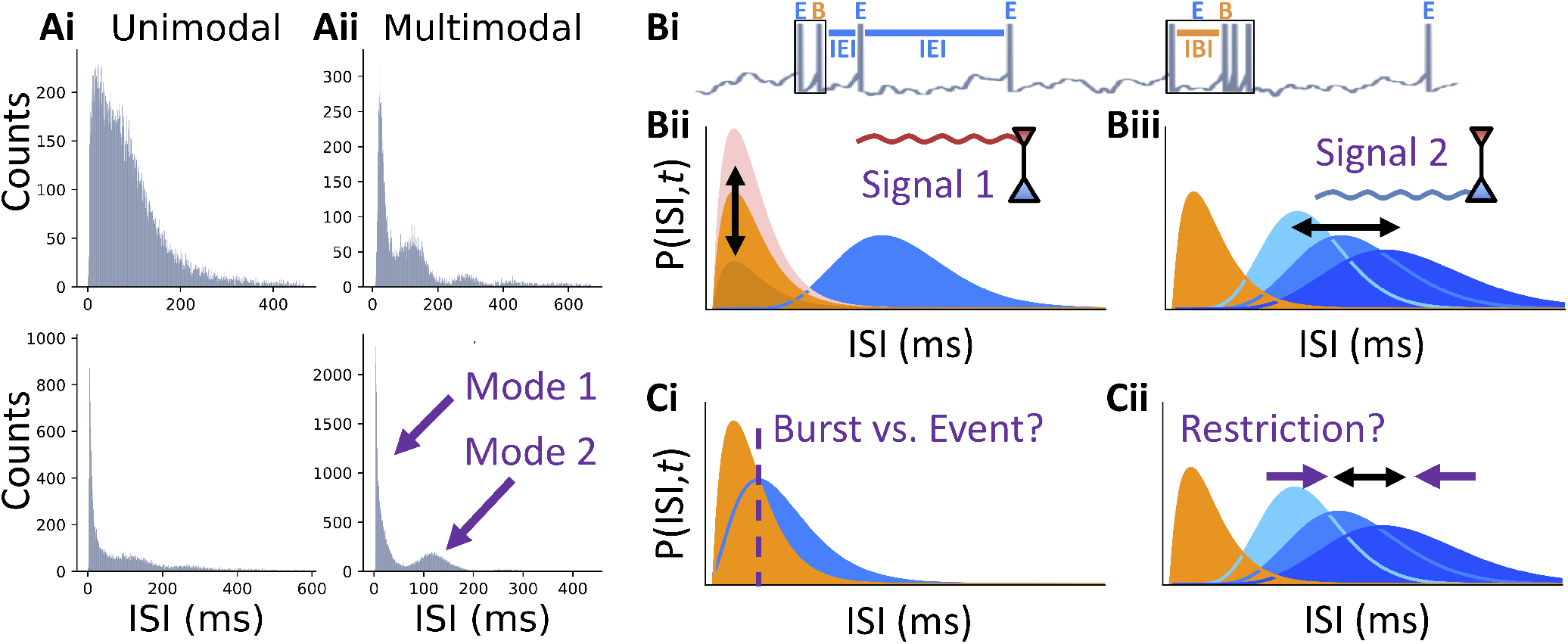
Inter-spike interval distribution and burst coding. **A**. Experimentally recorded Inter-Spike Interval (ISI) distributions have variable shape. Example ISIs from mouse visual cortex in vivo (first published in Ref. (1)), two showing a unimodal profile (left) and two showing multiple modes (right). **B**. Burst multiplexing decomposes spike trains into bursts (black rectangles labeled with orange B) and singlet spikes on the basis of the ISI widths. We then refer to any spiking event, either burst or singlet, as an event (labeled with blue E). ISIs within a burst are referred to as Intra-Burst Intervals (IBIs), e.g. the first ISI in the second burst of the pictured spiketrain, while those between events are denoted Intra-Event Intervals (IEIs), e.g. 2nd and 3rd ISIs of the spiketrain **(i)**. Burst multiplexing assumes that the instantaneous ISI distribution, denoted as a function of time by *P* (ISI, *t*), is a mixture of two components: a burst component whose amplitude encodes the first multiplexed signal (signal 1 impinging on inset neuron) **(ii)** and an event component whose mean encodes the second signal (signal 2 impinging on inset neuron) **(iii). C**. Burst multiplexing is impaired when the two components are overlapping because burst and event spikes become indistinguishable on the basis of the ISI **(i)**. Restricting the event component to bimodal ISI distributions may reduce the dynamic range for event rate **(ii)**.

Here, we quantified the linearly decodable information between inputs applied to a simulated ensemble of cells, utilizing the burst multiplexing (20) burst code (see Fig. 1Bi), and readouts that mimic synaptic processing as a function of changing firing statistics. Our results show that, while enforcing a bimodal ISI distribution does not appreciably affect information transmission, a burst code can still be implemented by cells with unimodal ISI histograms. Given this inability of the ISI histogram to distinguish burst-coding cell models from non-burst coders, we next asked whether other spike train statistics used for quantifying the propensity of a neuron to fire bursts of spikes (3, 23, 28, 29), might be equally ineffective at recognizing burst coding. We found that these metrics did not reliably distinguish burst-coding neuron models from non-burst-coding models, suggesting a disconnect between *visibly* bursty cells (*e*.*g*., with bimodal ISI distributions) and *functionally* bursty cells (*i*.*e*., those utilizing a neural code that attributes a particular meaning to bursts). Overall, our work details a rationale for using decoding approaches that separate bursts and isolated spikes even when a cell is not visibly bursty. Such approaches may reveal previously undetected streams of information within spiking data.

## Results

We investigated the relationship between the shape of the ISI distribution and information transmission by calculating information transmission between two simulated neuronal populations while varying the properties of the network. Our modeling approach is based on a cortical microcircuit where two streams of information are impinging on two distinct compartments of a population of pyramidal cells able to produce bursts, and to route these spike timing patterns to different post-synaptic neuron populations via target-specific short-term plasticity (Fig. 2A). We considered an *encoding* population (Fig. 2B), which received two inputs and explicitly emitted two types of *events*: single spikes or bursts. One of the inputs controlled the rate of events randomly generated amidst a relative refractory period. The other input modulated the probability that an event is a burst. When the model emitted a burst, it would add spikes to the spike train by sampling from a fixed distribution of short ISIs, the Intra-Burst Interval (IBI) distribution. Importantly, the encoding population did not force bursts to have a higher frequency than the highest achievable frequency of events. Instead, the IBI distribution could overlap with the Inter-Event Interval (IEI) distribution. We called this model the Burst-Spike-Response Model (BSRM) (see Methods) as it extends the spike-response model (30). As a reference to the theory of modulator-driver inputs (31) we called the input controlling the rate of event generation the *driver* and the input controlling the burst probability the *modulator*.

**Fig. 2.**
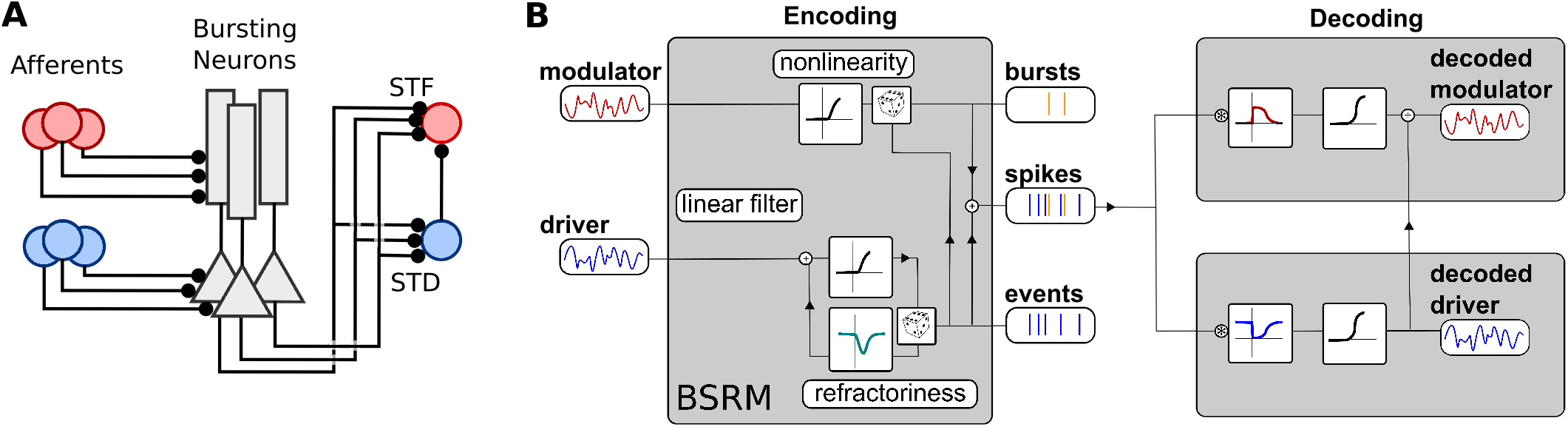
Schematic illustration of the simulation paradigm. **A** We model a population of bursting neurons receiving inputs from two distinct pre-synaptic populations and projecting to two post-synaptic cells with different short-term plasticity (short-term depression (STD); short-term facilitation (STF)). **B** Each bursting neuron is modelled with a Burst Spike Response Model (BSRM), which utilises distinct inputs to generate a bursty spike train. The driver input (blue, left) controls the event train, which is generated stochastically (illustrated by the die in the lower part) with a firing intensity that is a nonlinear readout of the input. A modulator input (red, left) controls the burst probability via another nonlinear readout. Upon event generation, a burst can be generated according to a Bernoulli process (upper die). Spike trains from this encoding cell population contain elements of both inputs. Two decoding neurons attempt to demultiplex the original signals. Different properties of short-term plasticity (illustrated by different linear filters before a nonlinear readout) extract bursts and events. A division from the event decoding cell is included to turn the estimate of the burst rate into an estimate of the burst probability, an essential step for extracting the modulator input.

Next we considered two neurons that were post-synaptic to the encoding population and attempted to retrieve both the driver and modulator inputs without knowing which spikes were generated from which distribution. These *decoding* neurons attempted to retrieve the time-dependent signals of both the modulator and driver inputs from the output of a single, uniform population. This demultiplexing was done by allowing the two post-synaptic neurons to have different types of Short-Term Plasticity (STP), whereby the efficacy of transmission depended on the preceding ISIs. For simplicity, we modelled STP by introducing an all-or-none dependence between the amplitude of the post-synaptic potential and the previous ISI. The driver-decoding neuron was excited by a spike from the encoding population only if it was preceded by an ISI above a fixed threshold, thus an extreme manifestation of short-term depression. Conversely, the modulator-decoding neuron received an idealized form of short-term facilitating synapses. These connections enacted the effect of a pre-synaptic spike only if it was preceded by an ISI below the fixed threshold. By previous work (20), the driver signal was expected to be decoded by the membrane potential of the downstream cell with depressing synapses. Conversely, the neuron with facilitating synapses retrieved the burst rate, a nonlinear mixture of driver and modulator signals. To decode the modulator signal, it was necessary to take the fraction of the burst rate and the event rate, an operation that could be implemented by divisive inhibition (20, 32). Here, we calculated the quotient of the two decoding cells’ membrane potentials. Information transmission was then quantified using the linearly decodable Shannon’s mutual information rate (33–35) (see Methods) by comparing the modulator and driver inputs to the cell potential quotient and the potential of the driver-decoding neuron. We refer to the information communicated between the modulator input and quotient of cell potentials as the modulator, or burst, channel, and the analogous driver-related quantity as the driver, or event, channel.

### Potent information transmission with unimodal inter-spike interval distributions: liminal burst coding

To manipulate the shape of the ISI distribution, we varied the parameter for the relative refractory period of events, *τ*_rel_. Small to medium values of the relative refractory period allowed very small IEIs such that the IEI and the IBI distributions overlapped (Figure 3Ai-ii). For short relative refractory periods, the overall ISI distribution was unimodal and the resulting spike trains had the appearance of a Poisson process (*i*.*e*. an exponential distribution). Large values of the relative refractory period were associated with events that were farther apart (Fig. 3Aiii). Since these could be either singlets or bursts, it resulted in sporadic bursts of spikes, a regime that should be ideal for burst multiplexing (20). When assessing how changing the relative refractory period affected information transmission, we considered two conditions and two decoders. In the *constant rate* condition, the cell firing threshold was adjusted to compensate for the changes in firing rate incurred by modifying the relative refractory period. Alternatively, we also simulated responses where we only varied the relative refractory period and allowed the firing rate to change accordingly. This setup will be referred to as the *uncompensated* condition. Since information loss is likely to arise from a misclassification of burst and singlets, we also considered a *perfect decoder*. The perfect decoder did not rely on the ISI for burst identification, but was given the information of whether a spike was generated by the driver or modulator input. Together, these different conditions and decoding approaches allowed us to determine how burst coding depends on the bimodality of the ISI distribution.

**Fig. 3.**
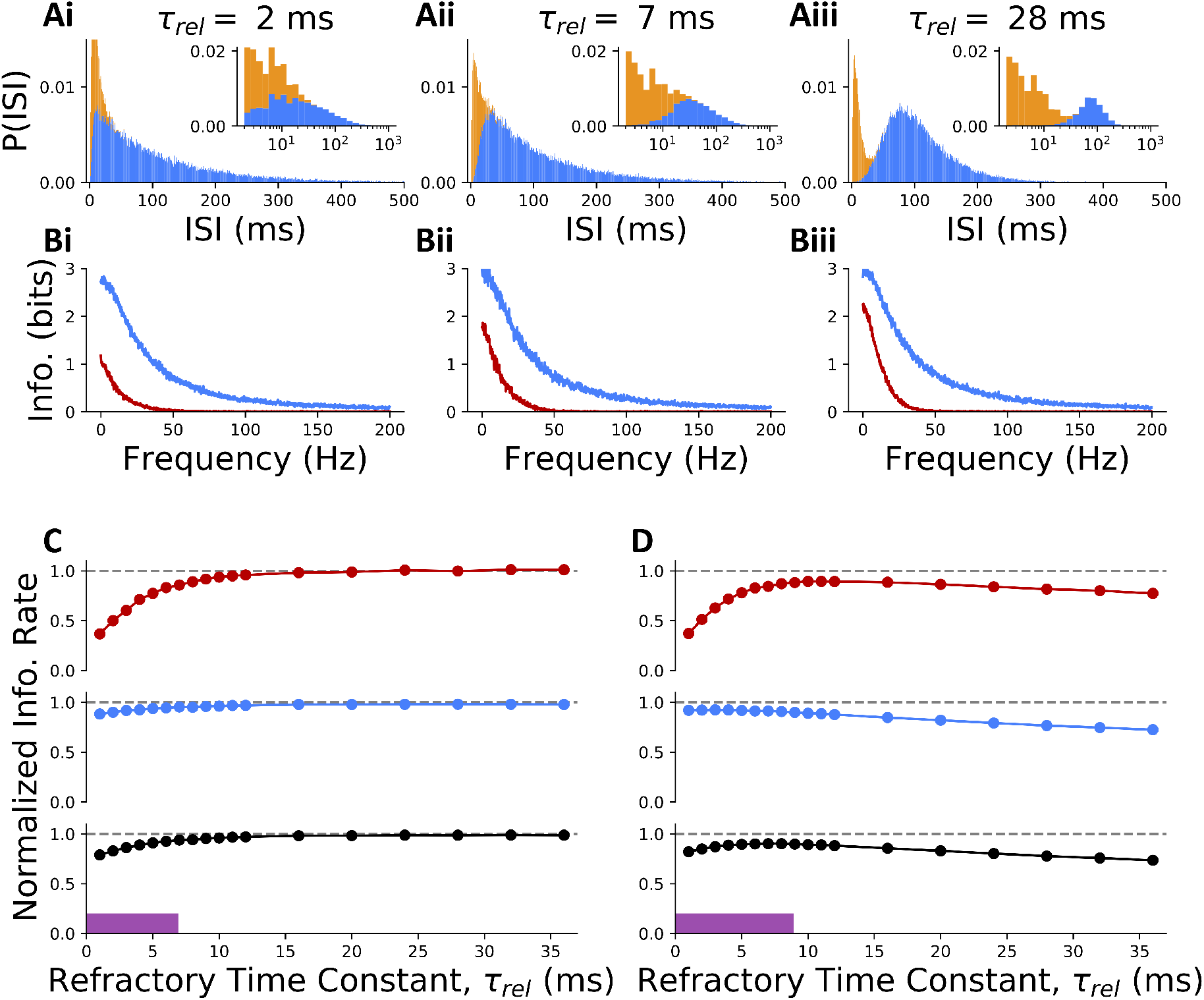
Burst coding despite unimodal inter-spike interval distributions. **A** The ISI distribution of the encoding population is plotted for three example relative refractory periods. Distribution is unimodal for very short *τ*_rel_ = 2 **(i)** and short *τ*_rel_ = 7 **(ii)** refractory periods, but bimodal for long *τ*_rel_ = 28 **(iii)** ones (rate compensated data is shown). **B** A lower bound on mutual information is plotted as a function of Fourier-frequency for both the modulator (orange) and driver (blue) channels. Integrating the lower-bounded information over frequency and summing driver and modulator channels produces estimates of linearly encoded information of **(i)** 131.58 bits/s **(ii)** 148.91 bits/s, and **(iii)** 156.64 bits/s for each distribution shown in (A), respectively. **C** Normalized Information rate is shown as a function of relative refractory period for the modulator channel (orange), the driver channel (blue) and both channels together (black). The purple bar indicates the region of parameter space producing visually unimodal ISI distributions. For each value of the relative refractory period, the firing threshold for neurons in the encoding population was scaled to preserve the same average event rate for all values of the relative refractory period. **D** As in C but without adjusting cell firing threshold. Normalization in C and D is by the maximum (over *τ*_rel_) linearly decoded information rate, for perfectly decoded events and bursts.

The linearly-decoded information spectrum between the modulator signal and the modulator-decoding neuron showed high information for slow input frequencies, which quickly dropped to zero for rapidly-changing inputs (Fig. 3B - red lines). For the driver signal, a low-pass profile is also observed, but with generally more information and a higher cut-off frequency (Fig. 3B - blue lines). These observations are exactly what is expected from the theoretical properties of the ensemble-burst code (20), whereby burst coding is optimal for the communication of more slowly changing inputs. In addition, the low stationary burst probability (here ≈ 0.2, chosen to match observations in cortex (9, 17)) is expected to reduce the proportion of information transmitted by bursts. To arrive at a more global measure of information transmission efficacy, we integrated the information spectrum over the linearly decoded information at each frequency (35), resulting in a measure of information rate in bits/s.

Next we focused on the total burst multiplexing information, calculated as the sum of the information rates of the modulator and driver channels (see Methods). In the constant rate condition, we found that both the modulator and driver channels were best transmitted as the ISI distribution of the encoding population became more bimodal. In this limit, both channels matched the perfect decoder (dashed lines in Fig. 3C-D). The driver channel was almost unaffected by the changing ISI interval distribution, transmitting 90.13% of the information even when almost the entire IEI distribution overlaped with the the IBI distribution (see Fig. 3Ai), with a 2 ms relative refractory period for events. Showing a drop to 50.10% in information at *τ*_rel_ = 2 ms, the modulator channel was more profoundly affected by using a short refractory time constant. Information started to degrade weakly just before the transition from bimodal to unimodal ISI distributions (at a relative refractory period of 7 ms), but the impediment remained weak when the ISI distribution became unimodal (purple bar in Fig.3C). A stronger transition to low information transmission occurred at a relative refractory period of 5 ms. As the driver contributes more information than the modulator channel, the total burst multiplexing information observed only a moderate decrease in information transmission (Fig. 3C black), and this at the highest overlap between distribution modes. We refer to the range where ISI distribution is unimodal, but the IEI and IBI show minimal overlap as the *liminal* regime. In this regime the IBI and IEI distribution join but hardly overlap, and thus bursts/non-burst can be separated reliably. We also note that an important fraction of modulator information (40-60% of perfect decoder) could be transmitted even when the IBI and IEI distribution entirely overlapped. We have reproduced these results (Fig. 3) using another related information theoretic approach (correlation theory, see Supplementary Note A and Supplementary Figure 8). Together, this highlights the possibility for cells without a clear demarcation between bursts and single spikes to use burst coding almost as effectively as cells with perfectly separated firing patterns.

In the uncompensated condition (Fig.3D), both the driver and modulator information were decreased with increasing bimodality. This reflects the effect of decreasing firing rate caused by increasing relative refractory periods, which naturally affects the information transmission. As a consequence, this condition is associated with a maximum of the modulator information over the the range of relative refractory period tested, a maximum that occurs in the liminal regime.

### Liminal burst coding is influenced by the properties of dynamic synapses

Next, we revisited some of the assumptions made on the properties of synaptic dynamics enacting the decoding. In particular, we were interested to know whether liminal burst coding could also take place when synaptic transmission was a smooth function of firing frequency – as observed in experiments – instead of the all-or-none dependence assumed in the previous section. To control the sensitivity on firing frequency, we varied the function relating synaptic efficacy and previous interspike intervals. Using sigmoidal relationships, we could interpolate between a sharp, (Fig. 4Ai-ii) and a smooth (Fig.4Bi-ii) dependence of synaptic efficacy on the previous interspike interval. It was not immediately clear to us if a more graded dependency would be beneficial to the information transmission. Burst coding could be perturbed by a graded dependency, as each of the variations in ISIs are transmitted as fluctuations in post-synaptic potential amplitudes. But it could be beneficial because a graded relationship between ISI and efficacy could implement a concept of confidence in the categorization. Figure 4C shows results for the modulator signal only, as it is more strongly corrupted by unimodal ISI statistics, although the sensitivity to firing frequency was modulated in both burst and event channels. Decreasing the sensitivity of the decoding synapses to firing frequencies (i.e. increasing the smoothness) made the drop in information due to bimodality more graded. In addition, decreasing the sensitivity to firing frequency tended to reduce burst coding efficacy (Figure 4Biv). These results demonstrate that burst transmission is reduced slightly when synapses have graded sensitivity to firing frequency; however, this reduction is negligible when the graded sensitivities are appropriately steep.

**Fig. 4.**
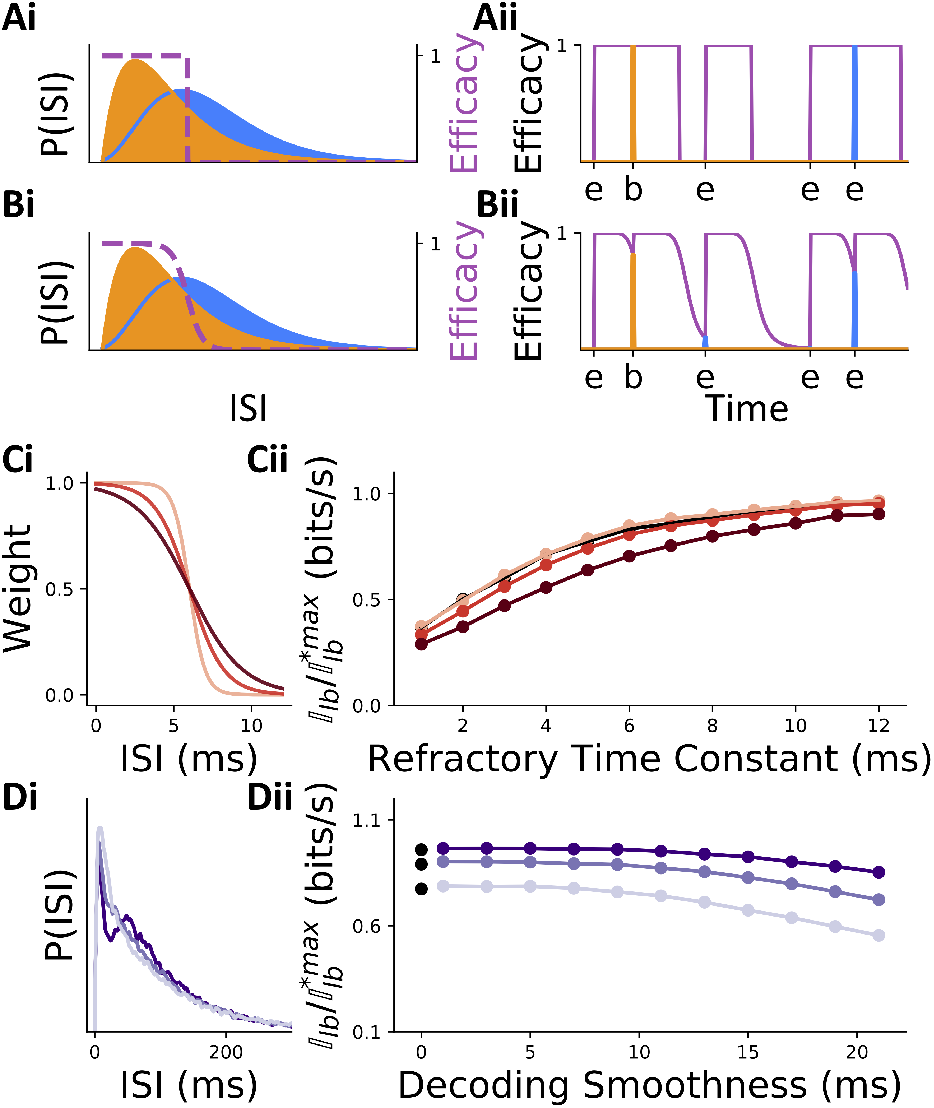
Properties of Dynamic Synapses Influence Information Transmission. **A-B** Bursts and events can be decoded by either sharp (A) or smooth (B) dependence of the synaptic efficacy on the interspike interval. IBI (orange) and IEI (blue) distributions are overlayed on the function relating interspike intervals to synaptic efficacy (purple, right axis). **Aii-Bii** The effect of this function on the decoded modulator signal is illustrated using a single hypothetical spike train with true events (ticks labeled by ‘e’) and true intra-burst spikes (ticks labeled by ‘b’). The strength at which an incoming spike will be read out is represented by the height of the blue and orange vertical lines (event = blue; intraburst = orange). **Ci** Three instances of graded sensitivity to the firing frequency. **Cii** Modulator information transmission is shown against the refractory time constants around the liminal regime for the three nonlinear sensitivity functions shown in Bi (same colour scheme – rate compensated data shown). Black line (obscured behind yellow) gives sharp threshold-based communication for comparison. **Di** ISI distributions for three different relative refractory period time constants: 5 ms (light purple), 8 ms (purple), 12 ms (dark purple). **Dii** Modulator information transmission as a function of the sensitivity to firing frequency for the three ISI distributions pictured in (iii) (same colour scheme). Black dots denote sharp threshold-based communication. All data for variable *τ*_rel_ is rate corrected, as in Fig. 3C, and data for sharp threshold communication is taken from Fig.3.

### Burst length and low event rates

We next considered other parameters that contribute to the efficacy of burst coding. The first parameter we investigated was burst length. We therefore varied the number of spikes following the first spike in a burst, namely the number of intra-burst (IB) spikes. The burst multiplexing code used here differentiates only between bursts and singlets and is therefore agnostic to the number of IB spikes. If burst length is increased from doublets to longer bursts of *N* IB spikes, the STP rule of the decoding cell can be adjusted (see Methods section) to respond only to spikes after being primed by *N* − 1 IBIs (Fig. 5.Ai). Such a dependence between the number of intra-burst spikes and the changes in synaptic efficacy is observed in experiments (36). Since the probability of observing *N* subsequent events with short ISIs decreases as *N* increases this allows post-synaptic cells to better distinguish events from bursts, regardless of the overlap between IEI and IBI interval distributions.

**Fig. 5.**
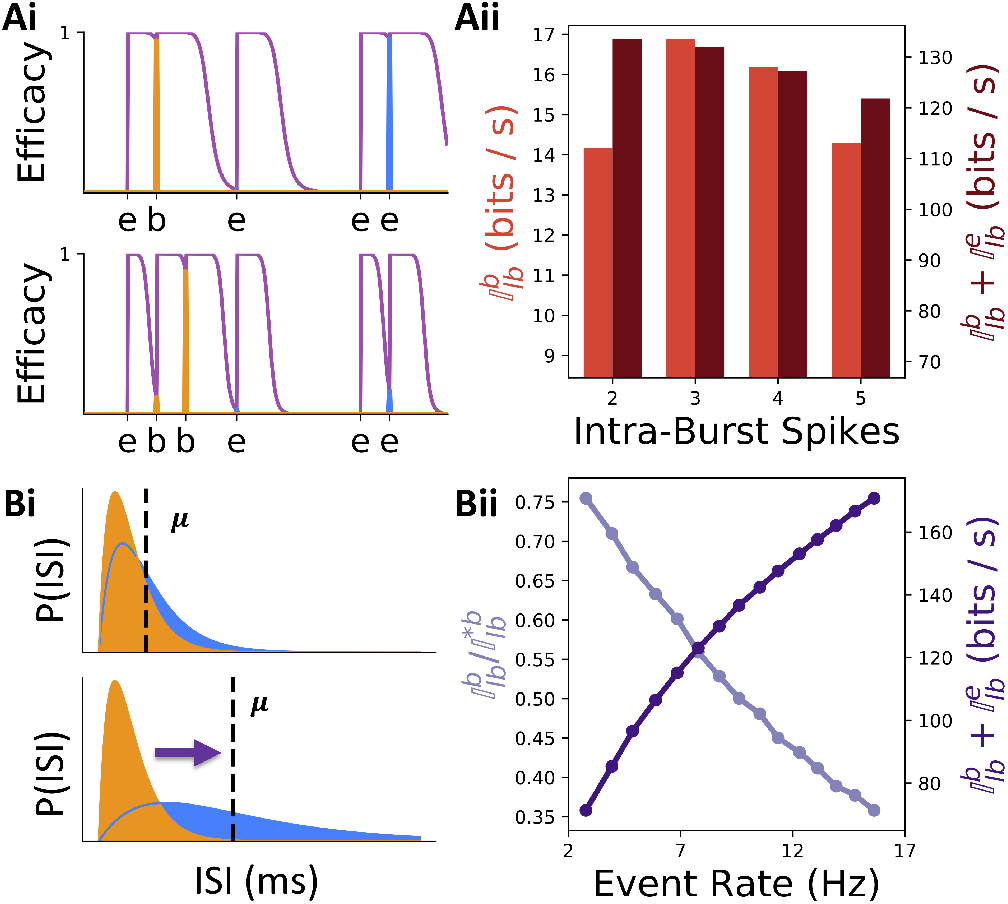
Effects of Burst Length and Event Rate on Information Transmission. **Ai** Illustration of the modulator-channel communication of a single hypothetical spike train made of a number of true events (‘e’ ticks) and bursts (‘b’ ticks) with IBIs comparable to IEIs (as in Fig. 4Aii and Bii). **Top:** for bursts made of 2 IB spikes, to communicate the burst implies that an event with a similar interval (last event) will also be communicated. **Bottom:** using bursts made of 3 IB spikes (orange) allows the burst to be transmitted while minimizing transmission from the last event because the synaptic efficacy is allowed to accumulate through successive IBIs. **Aii** Modulator channel information rate (left y-axis) and total information rate (right y-axis) are shown as a function of the number of intra-burst spikes. **Bi** Schematic illustration of how the overlap between a fixed IBI distribution (orange) and the IEI distribution depends on whether the average IEI (*µ*, dashed line) is short (top) or long (bottom). **Bii** Normalized burst channel decoding efficacy (left y-axis) monotonically decreases, with increasing average event rate, while the total burst multiplexing information (right y-axis) increases. Both Aii and Bii were generated using strongly unimodal ISI distributions (*τ*_rel_ = 2 ms.)

We found that increasing burst length increased burst channel information, particularly for 3-4 IB spikes (Fig. 5Aii). The total information, however, always decreased with increasing burst lengths. We know of two factors limiting information transmission as burst length is increasing. First, there is a poorer temporal alignment between the burst generating signal and the transmission of bursts due to an increased number of randomly sized intra-burst ISIs between burst generation and burst detection. This error could, in theory, be reduced in cells with less randomness in their IBIs. Second, though they improve burst transmission, longer bursts do not contribute any new information to a spiketrain but still occupy significant periods of time, causing fewer event spikes to be fired per unit time and thus decreasing information transmission rate for events. The latter effect was discussed by Naud and Sprekeler (2018) (20), where it was shown that increasing burst length is detrimental in general, but that study did not take into account the influence of synaptic properties on total information transmission. Our results supplement this theoretical study and indicate that increasing the length of bursts to more than two spikes per burst is beneficial only to the communication of a modulator signal.

Next we considered reducing the average event rate. In Fig. 3C, we had kept the average event rate fixed to a value close to 10 Hz. Reducing the average event rate results in a lower spike rate and thus lower information rate, but also reduces overlap between IEI and IBI distributions (Fig. 5Bi), thus limiting spike misclassification error and improving burst channel efficacy. To quantify the relative improvement in burst-decoding as event rate is decreased, we performed simulations using clearly unimodal ISI distributions (*τ*_rel_ = 2 ms) and varied the average IEI. To quantify information transmission amidst changing firing rate, we normalized the constantrate information transmission, 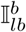, by the information calculated with perfect decoding 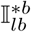. For the transmission of modulator information, we found that transmission improves as event rate decreases (Figure 5Bii - light purple). Concomitantly, the total information rate decreases with decreasing event rate (Figure 5Bii - dark purple). Together, we found that the average event rate controls a compromise between the communication of the modulator input and the communication of the net amount of information even when IBI and IEI distributions overlapped.

### Classic methods fail to identify functional burstiness

A large body of research has been devoted to distinguishing cells that burst from non-bursty cells (12, 37–39). Some of the methods developed to this end (23, 26, 28, 29) have focused on classifying cells based on the similarity of their response statistics to those of the Poisson process. Other approaches have focused on the identification of separate peaks in the ISI distribution or the auto-correlation function. Given that a population of neurons can communicate burst-coded information efficiently despite a unimodal ISI distribution, we now ask whether this disconnect between the bimodality of the ISI distribution and the functional role of bursts extends to other metrics for classifying bursty cells.

We first compared the ISI distribution and autocorrelation of a Poisson process with absolute refractory period (23) to those of our BSRM model. Figure 6A-B shows one example where the BSRM model generates an ISI distribution and autocorrelation that were strikingly similar to the Poisson process. Since the BSRM is explicitly coding information in bursts but the Poisson model is not, these results show that any metric based on the ISI distribution or the autocorrelation function will not always detect burst coding.

**Fig. 6.**
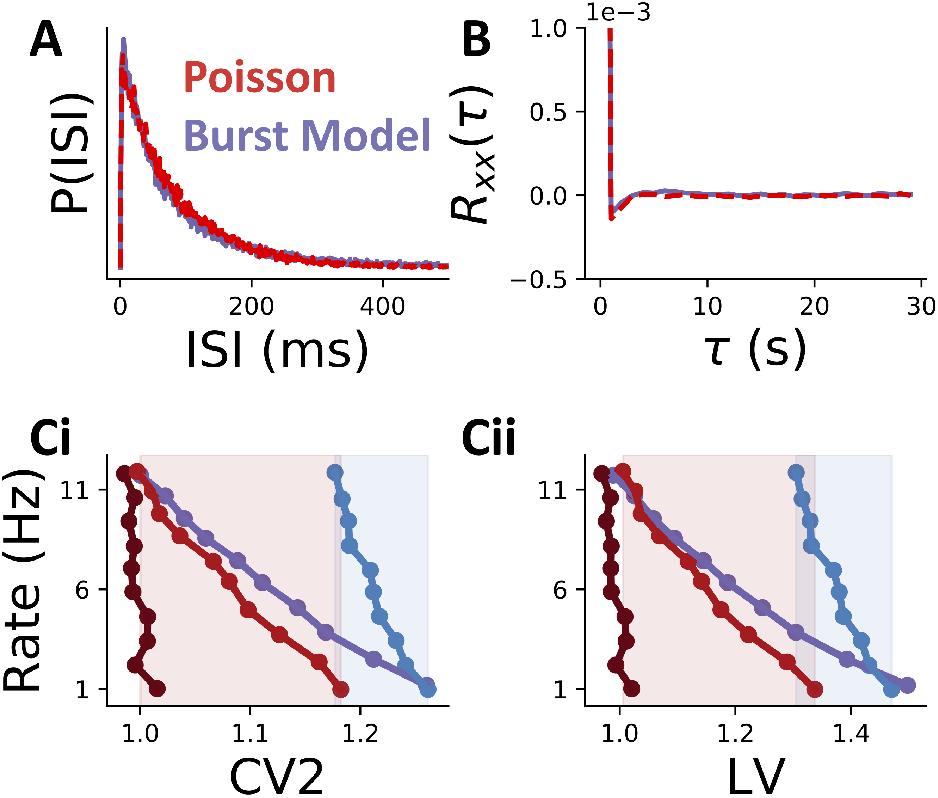
Common metrics of spike train irregularity do not distinguish functionally bursty cells. **A** ISI distribution of the BSRM model in effective multiplexing regime (solid, purple) is almost identical to that of a homogeneous, refractory, Poisson process (dashed, red). **B** Autocorrelation of spike trains from (A) are almost identical. Y-axis units are spikes^2^. **C** Plotted are values of a given spiketrain statistic (x-axis) for a range of neuron spike rates (y-axis) for three different models: a single-input rate model with high input signal variance (light red), the same rate model with low input variance (dark red), a stationary gamma renewal process with shape parameter 0.5 (blue), and the BSRM with 6 ms refractory period (purple). LV **(i)** and CV2 **(ii)** values of the BSRM overlap substantially with the high-variance input rate coding model (red shaded region) and stationary gamma renewal process (blue shaded region). All data points in this figure were calculated from simulations of 4800-second long spiketrains, with a 1 ms discretization.

Next, we tested whether previously used metrics of spike-train irregularity, the Coefficient of Variation 2 (CV2; Fig. 6Ci) (28) and the Local Variation (LV; Fig. 6Cii) (29), were able to distinguish burst-multiplexing from non-burst-coding models. Both of these statistics take into account the relative order of ISIs in an effort to quantify the “local” variability in a spiketrain–that is, how similar adjacent ISIs are. This makes them more robust to changes in firing rate than, for example, the coefficient of variation of the ISI distribution. Intuitively, this might allow them to pick up on the added burst structure of the BSRM. To test this idea, we simulated the BSRM along with two non-bursting neuron models and checked whether the models could be parameterized to yield statistics indistinguishable from the BSRM. We found that both CV2 and LV of a Spike-Response Model (SRM; (40)) neuron communicating a single input without a special meaning for bursts were distinct from the CV2 and LV of a BSRM only when the SRM received inputs with low variance. When the SRM received inputs with high variance, both the CV2 and LV were larger, covering almost the entire range of values produced by the BSRM. We found that CV2 *>* 1.2 and LV2 *>* 1.4 were not easily generated by the univariate SRM, although these values were generated by the BSRM at a low firing rate. We then asked if another non-burst coding neuron model would produce the CV2 and LV values in this range. We found that the stationary gamma renewal process – which approximates the firing statistics of the leaky integrate and fire model (41) – covered this range of values of CV2 and LV (blue lines in Fig. 6C). Together, the range of CV2 and LV values obtained from the BSRM while using bursts to communicate to streams of information is covered by realistic models of firing that do not utilize burst coding. Overall, we could not find a single-spiketrain metric that was able to reliably recognize burst coding at work.

## Discussion

This paper has, primarily, contributed two findings: (i) Uni-modal ISI distributions do not preclude burst coding in the form of burst multiplexing; (ii) classic metrics for identifying bursty cells are unable to recognize when burst coding is being utilized. Here, we discuss the mechanisms and implications of these findings.

The reason measures of local variation are unable to fully separate the BSRM from rate models is that the input signal to a rate model can induce highly irregular spiking in the model if it combines high power, for large fluctuations, with a slow fluctuation time scale, resulting in short periods of supra-threshold input and fast, burst-like firing, followed by periods of semi-quiescence with long ISIs, when the signal fluctuations fall below threshold. Conversely, rapidly fluctuating input signals tend to result in less irregular spiketrains, resulting in CV2 and LV values close to that of a homogeneous Poisson process (dark red lines in Fig. 6C). These two different input signal regimes result in very different SRM ISI distributions, with the rapidly-changing signal exhibiting an ISI distribution akin to the Poisson process and the slowly fluctuating signal exhibiting an ISI distribution similar to the Gamma renewal process. While CV2 and LV alone were unable to separate BSRM and SRM, augmenting these statistics with extra information, such as cell ISI distribution and firing rate might aid in the classification of burst-coding cells. For example, if the ISI distribution is similar to a Poisson process, making spiketrain irregularity due to input signal unlikely, but LV or CV2 are high, this might be suggestive of a functionally bursty spiketrain.

The failure of classic statistics to identify burst multiplexing cells has two main implications. First, a larger class of cells could be utilizing bursts to code information than previous analyses have led us to believe. The robustness of burst multiplexing to the shape of ISI distributions but also to variants in synaptic decoding mechanisms suggests utility for the burst multiplexing code in biological and artificial systems. For the brain, this result implies that burst multiplexing can be implemented by more cell types and synaptic plasticity rules than one might previously have thought. As such, it becomes clear that one cannot dismiss the occurrence of burst multiplexing simply because a given cell displays unimodal ISI statistics. This has implications for the recently proposed theory that burst multiplexing is instrumental for the coordination of plasticity and for allowing biological networks to solve tasks that depend on hierarchical architectures (21). Thus, our work implies that cells that do not show prominent bursts may also utilize these spike-timing patterns to coordinate plasticity.

Second, our work motivates a distinction between two types of “bursty” cells: those that are functionally bursty, i.e. utilize bursts to code information, and cells that are visibly bursty, i.e. showing two modes or a clear peak in mass at low values in their ISI distributions. It seems likely then that the class of visibly bursty cells is simply a subset of the functionally bursty cell set, although it is also possible that some rate coding cells fire bursts that are not used for coding. If none of the metrics are likely to establish the presence of burst multiplexing in a spike train, then what will? Our work argues that a population approach is required. Targeted methods, such as targeted dimensionality reduction (42, 43), which could search for specific correlates in multi-dimensional patterns of activity could readily be applied to data *in vivo*. Our results argue that there is no ground to restrict the search for the correlates of burst coding to only those cells that appear to be bursting. Thus, we believe that this work provides grounds for heightened investigation of temporal codes in biological and artificial cell networks.

## Methods

### Model

The network model was composed of a population of two-compartment burst-spike response model (BSRM) cells receiving identical synaptic inputs (Fig. 2B), and two cells post-synaptic to this encoding population. The two inputs to the encoding population controlled the burst probability (modulator input) and the event rate (driver input) of its neurons.

#### Input signals

Ornstein-Uhlenbeck (O-U) processes simulated via the exact method (44) were used for the inputs, 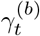 and 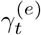. These were defined by

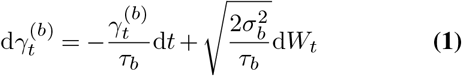

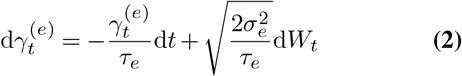

where d*W*_*t*_ is the Wiener process, *τ*_*x*_ is the time constant and 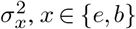 is its asymptotic variance. Higher variance and a larger time constant were used for the burst input as these were found to work better empirically. We expect the effect of the latter adjustment is because the variability inherent in intra-burst ISIs adds more noise to higher frequencies.

#### Encoding cell model

The BSRM is a modified version of the bursting rate model used in Naud & Sprekeler’s 2018 work (20). The difference is that we explicitly modelled intra-burst spikes. Specifically, BSRM is a self-inhibiting marked point process defined by the double sequence {(*B*_*n*_, 𝒯 _*n*_) | *n* ∈ *ℕ*} where 𝒯 _*n*_ is a process with rate *λ*_*t*_ and is constructed by alternating, as a function of the mark sequence, between sampling a modulated renewal process, for event spikes, and a renewal process, for intra-burst spikes. The rate is thus dependent on the mark process and is defined as follows

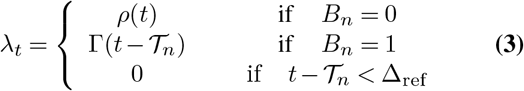

where *n* is the index of the last spike, *B*_*n*_ ∼ Bernoulli (*p*(𝒯_*n*_)) if the *n* − 1^*th*^ spike is not the first spike in a burst, in which case *B*_*n*_ is fixed to 0. *p*(*t*) is a function of the modulator input via the equations

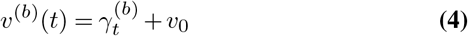

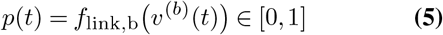

and *ρ*(*t*) is a function of the driver signal via

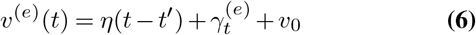

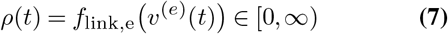

Finally, Γ(*t*) is the rate function for sampling an interval that is gamma distributed with scale parameter Γ_1_ and shape parameter Γ_2_ and *S*_*t*_ = Σ _*n*_ *δ*(*t* − 𝒯_*n*_) is the generated spike train, which is defined as a sum of Dirac delta functions over spike-time indices (40). The marked process labels the first spike in a burst with a 1 and all other spikes with 0.

The above model describes the doublet BSRM that was used in most of the results. For the BSRM with *N* intra-burst spikes, used in Fig. 5, we simply sampled *N* ISIs after the first spike in the burst from the gamma distribution. Thus, the only difference from the doublet model is that there are now *N* + 1 intra-burst spikes in a burst instead of just 2.

The following link functions were used to generate burst probability and event rate from the membrane potentials employed in the BSRM model.

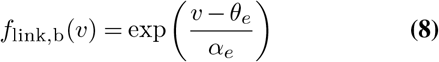

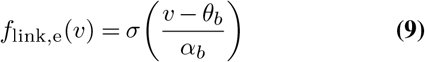

The exponential link function was chosen as it is commonly used in neural rate models (40). The sigmoid link function, denoted by *σ* above, was selected because it is the natural option for converting values on the positive real line to probabilities and has been used for burst modeling historically (20). In both functions *θ*_*x*_ represents a threshold parameter and *α*_*x*_ determines the sensitivity of the threshold (*x* ∈ {*e, b*}).

An exponential function was used to model the relative refractory time period of the neurons, as is standard (40), because the exponential decay reproduces the biological phenomenon. Using *t*^*′*^ as the time of the last spike:

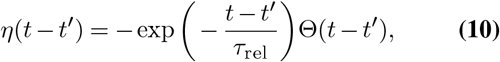

where Θ(*x*) is the Heaviside function with Θ(*x*) = 1 if *x >* 0 and 0 elsewhere.

We will now outline our rationale for parameter choice in the BSRM model. The cell resting potential doesn’t effect information transmitted and was set to zero for simplicity. The burst threshold and scale factor were chosen so that the fraction of total events that are bursts was approximately 0.2, as the burst fraction measured in layer 2/3 and layer 5 cortical cells is measured to be in the range 0.1 to 0.2 (9). The burst ISI scale parameter was chosen so that rate of spikes generated by a sequence of burst ISIs would be in the 100-200 Hz range observed in layer 5 cells (45) and the shape parameter was selected to qualitatively produce the sharp, super-exponential peak observed in ISI distributions (46). The 2 ms absolute refractory period is in line with the literature on cortical cells (47) as well as cells in sub-cortical regions (48). Firing rate in cortical cells covers roughly two orders of magnitude, from around 1Hz to tens of Hz (49). Accordingly, the event threshold and scale factor were set to produce an event rate of approximately 10 Hz when *τ*_rel_ was set to 6 ms, which led to a value of 3.29 for the event threshold. For the uncorrected rate results (Figure 3.D) these initial event threshold and scale factor values remained fixed as *τ*_rel_ increased while for the rate corrected results the event threshold was decreased with increasing *τ*_rel_ to keep event rate fixed. Lastly, 200 BRSM cells were used for the encoding population as the goal was to explore a regime where spike generation (finite size effect) noise would be appreciable.

#### Decoding cells

For the decoding cells, we adopted the model described in Ref. (36), that defines STP weight functions as a composition of a linear filtering and nonlinear function. This can be formalized as

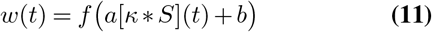

where *f* : ℝ → [0, 1] is a nonlinear function, *κ*(*t*) is a convolution kernel that we call the *weight function* because it encapsulates the frequency dependence of the synapse. The symbol ‘*’ denotes the convolution operation and *a* and *b* are parameters that are used to determine whether STP is facilitating or depressing. For sharp frequency dependence we set *f* (*x*) = Θ(*x*) and *κ*(*t*) = Θ(*t*) − Θ(*t* −*θ*_w_) and *θ*_w_ is the ISI threshold parameter, above which spikes are considered event-related and below which they are considered intra-burst. We will refer to this decoding model as STP1 and we note that it weights each spike as a function solely of the ISI that came directly before it (renewal dynamics).

We also designed a STP rule with a smoother dependence of the weight function on ISI and a dependence on firing history beyond the just-preceding ISI (as used in Figs. 4 and 5B). To achieve these desiderata, we chose *f* (*x*) = *σ*(*x*) and 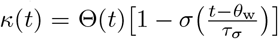, where *σ* denotes the sigmoid function and *τ*_*σ*_ is a parameter to control the smoothness of the function. We will refer to this model as STP2.

The above weight function definitions were inserted into the following equations which map spike trains of an encoding population of *N* cells to decoding-cell membrane potentials

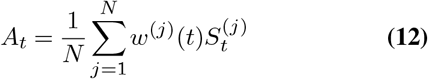

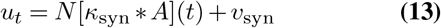

In these equations, *A*_*t*_ is the mean of weighted spike trains from the encoding population and *u*_*t*_ is the synaptic response of the post-synaptic cell. In this way, the event rate is estimated in the membrane potential of a downstream cell employing depressing STP, *u*^(*e*)^. To extract burst fraction, we must divide the burst rate by the event rate, an operation that has been shown to be implementable by neural machinery (i.e. divisive inhibition (20, 32)). Rather than explicitly modeling such neural machinery, we simply took the ratio of the estimated burst and event rates 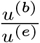, where *u*^(*b*)^ is the membrane potential of a neuron with short-term facilitation and estimating the afferent burst rate.

We found that information was better transmitted if a lag was introduced between *u*^(*e*)^ and *u*^(*b*)^ before taking their quotient. This makes sense given that the burst spike train lags behind the event spike train by the length of the intra-burst ISIs.

The synaptic filter *κ*_syn_ was modelled as an exponential rise and exponential decay:

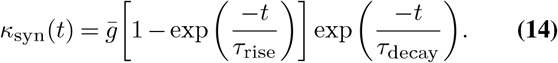

where 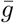 sets the scale of the synaptic conductivity.

The important aspect of the decoding cell parameterization was to arrive at quantities which were equivalent, in an information theoretic sense, to the synapses of decoding cells. For this reason their values were somewhat arbitrary, baring the following considerations. The resting membrane potential does not affect information transmission but, if set to zero, would result in an undefined estimate of burst fraction in a period of prolonged encoding population inactivity. The synaptic filter scale doesn’t effect information and was set for simplicity. Because the synaptic filter itself is linear it will not effect information and was included simply for completeness.

To set the lag between burst and event signals in the decoded burst fraction for all experiments with doublet spikes, we performed a line search to maximize decoded information on lags between 0 and 15, with increments of 1ms. This was done for 3 BSRM cells with *τ*_rel_ values of 2 ms, 6 ms and 12 ms. Based on this we set the lag in all cases of doublet spikes to 9 ms.

To set the weight function thresholds we first assumed that there would be a single threshold, above which spikes would be considered event-related and below which they would be considered intra-burst spikes. This threshold, *θ*_w_, was selected uniquely for each cell model in the study by performing a line search on 5 to 35 ms (5 to 45 ms for figure 5.B), in 1 ms increments, to maximize linear-decoding information.

The last parameters were those associated with STP rule 2. In figure 4 *a* and *b*, the facilitation weight function parameters, were set to 40 and −20 respectively. In figure 5.A these were set to 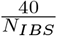 and −20, where *N*_*IBS*_ is the number of intra burst spikes modelled. In figure 4 *τ*_*σ*_ was varied in 2 ms increments between 1 and 20 while in figure 5.A it was set to 5 ms.

Finally, we note that, for STP1, equation 12 can be rewritten to match a previously used formalism (50) for describing weighted spiketrains

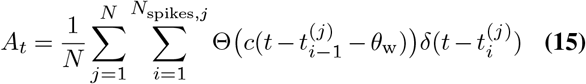

if, in equation 11, we set *f* to be the identity function, *a* = 1 (− 1), and *b* = 0 (1) for facilitation (depression). Here 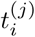 denotes the *i*th spike of the *j*th neuron, *N*_*spikes,j*_ denotes the number of spikes of the *j*^*th*^ neuron, *c* = −1 if the synapse is depressing and *c* = −1 when the synapse is facilitating.

### Estimating linear-decoding information rate

To quantify information in our system we employ information rate, a tool commonly used in theoretical neuroscience (8, 34). Naively estimating information rate is difficult because it involves the non-parametric estimation of mutual information between high dimensional vectors, a task that requires prohibitively large data sets (51). To make this estimation more tractable we rely on two methods that make use of the simple statistical structure of the Discrete Fourier Transform (DFT) of stationary Gaussian processes to estimate the linear-decoding information rate. The method used to estimate this rate throughout the results section is Stein’s method (35); in the supplementary section (see supplementary material) we provide novel insight into the recently proposed correlation theory method (52) and show that it reproduces our results with the former technique (see Figure 8).

**Fig. 7.**
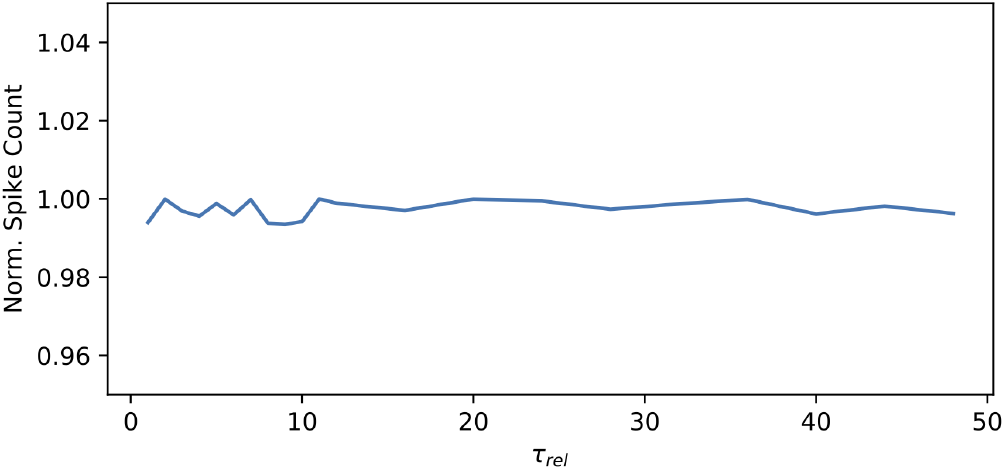
Figure 3 Graphical demonstration that rate correction in figure 3 was correctly implemented. Y-axis is the total spike counts per ISI distribution, normalized by the maximum of these over parameter values. X-axis is parameter value

**Fig. 8.**
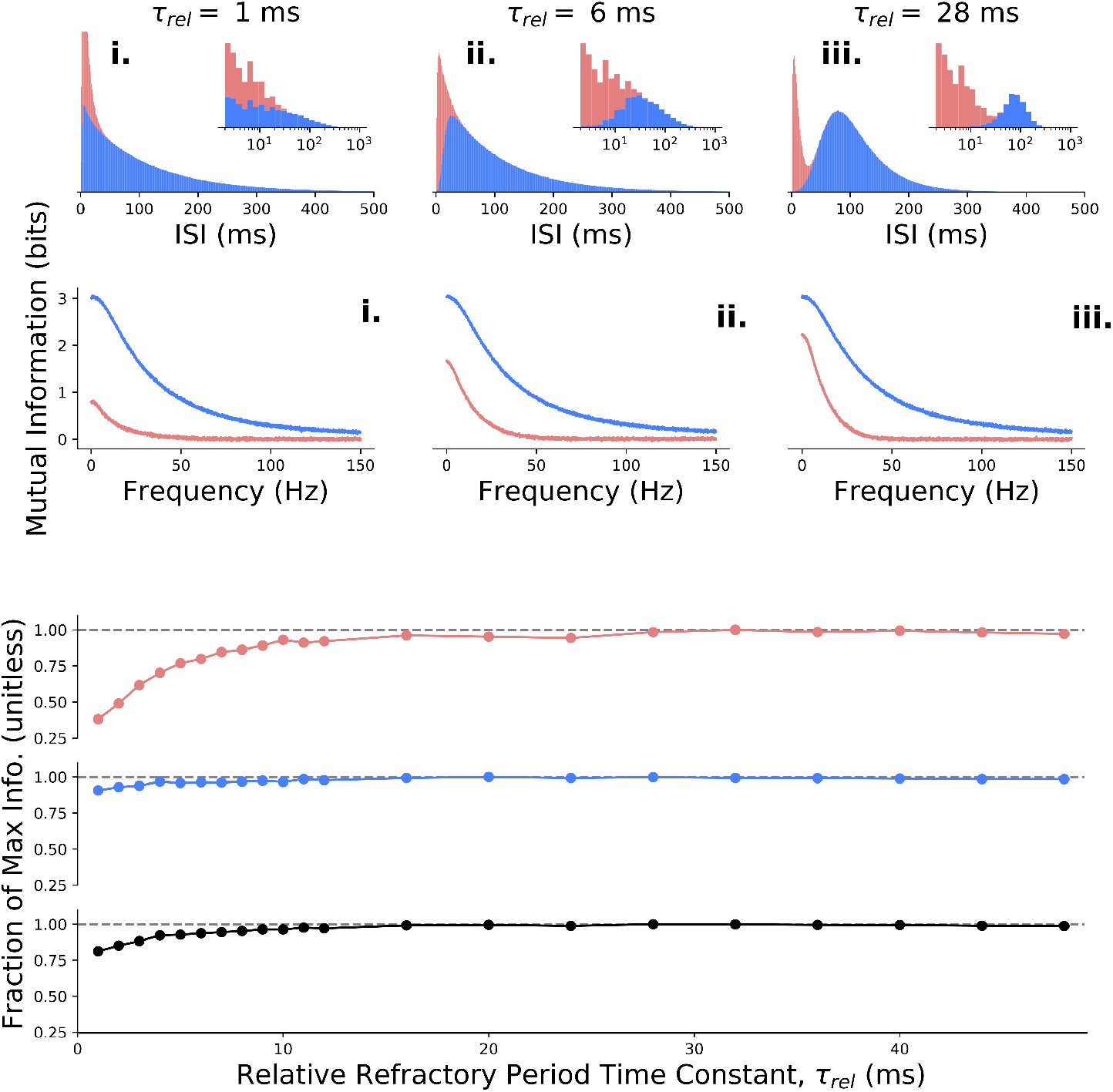
Figure 3 reproduced with correlation theory. Note that burst channel information is plotted in orange here rather than the red used in Fig. 3.

Stein’s method is given by the following equation (35):

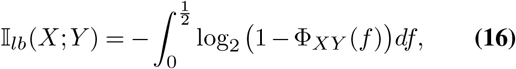

where Φ _*XY*_ denotes the coherence between *X* and *Y*, which must be estimated from the data. This method requires the input signals to be stationary Gaussian processes, constraints which were satisfied by our use of O-U processes as stimuli signals.

To implement Stein’s method we estimated the power spectra of input and output processes and the cross spectra of the two, then used these to calculate the coherence. Estimation was performed with Welch’s method (53), in Scipy, with a Hanning window. The parameters of this method are given in table 4. Here *T* is simulation length and *N*_*win*_ is window length. The overlap was chosen to be half the window length throughout. Simulations were run 5 times with different random seeds; all plots are the trial means.

**Table 1.**
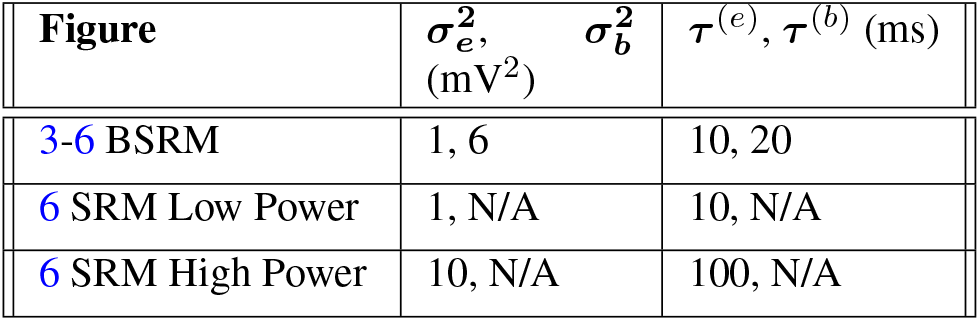
Power and timescale of the Ornstein-Uhlenbeck input signals.

**Table 2.** Model parameters for the encoding cells

| Name | Symbol<br>(Units) | Value |
| --- | --- | --- |
| Cell Resting Potential | $v_0$ (mV) | 0 |
| Burst Threshold | $\theta_b$ (mV) | 4.5 |
| Burst Scale Factor | $\alpha_b$ (mV) | $\frac{1}{3}$ |
| Burst ISI Distribution Scale | $\Gamma_1$ (ms) | $\frac{20}{3}$ |
| Burst ISI Distribution Shape | $\Gamma_2$ (unitless) | 1.5 |
| Absolute Refractory Period | $\Delta_{\text{ref}}$ (ms) | 2 |
| Relative Refractory Time Constant | $\tau_{\text{rel}}$ (ms) | Variable |
| Event Threshold | $\theta_e$ (mV) | Variable |
| Event Scale Factor | $\alpha_e$ (mV) | 2 |
| Encoding Population Size | $N$ (unitless) | 200 |

**Table 3.** Model parameters for the decoding cells.

| Name | Symbol (Units) | Parameter Value |
| --- | --- | --- |
| Resting Potential | $v_{syn}$ (mV) | 1 |
| Synaptic weight | $\bar{g}$ (mV) | 1 |
| Synaptic Rise Time | $\tau_{rise}$ (ms) | 3 |
| Synaptic Decay Time | $\tau_{decay}$ (ms) | 5 |
| STP1 Depression | $a, b$ (none) | -1, 0.5 |
| STP1 Facilitation | $a, b$ (none) | 1, -0.5 |
| STP2 Depression | $a, b$ (none) | 40, -20 |
| STP2 Facilitation | $a, b$ (none) | Variable |
| STP ISI Threshold | $\theta_w$ (ms) | Variable |
| STP Smoothness | $\tau_\sigma$ (ms) | Variable |

**Table 4.** Hyper-parameters for the estimation of information.

| Figure | $T$ (s) | $N_{win}$ (s) |
| --- | --- | --- |
| Figure 3, 5.Bii | 1007.616 | 8.192 |
| Figure 4, 5.Aii | 503.808 | 4.096 |

### On Summing Burst and Event Channel Information Rates

We used the sum of burst and event channel information rates, rather than the information rate between bivariate input and output signals. Because the bivariate information rate considers the full information between input and output vectors, it does not effectively quantify the extent that the inputs are “demixed” in the decoded outputs. For example, because mutual information is agnostic to invertible transformations, the bivariate method would assign equal information rates to a model whose decoded signals at a given time point are an invertible transformation of the input burst and event signals, at that time point, and a model that perfectly demixes the two inputs.

It was not immediately clear, however, whether the sum of mutual information rates would yield wrongly high rate estimates by counting information twice, as would occur if the input modulator and driver signals were precisely equal. This worry is easily resolved if one uses independent modulator and driver inputs, as in this study, which we prove below for completeness.

To prove that the sum of information rates does not count information twice, we show that the information rates for the burst and event channels is less than the information rate for the two channels combined when modulator and driver inputs are independent. We first make some definitions: the input and output stochastic processes are *𝒢* = {Γ_*t*_}_*t≥*1_ and U = {*U*_*t*_}_*t≥*1_ respectively, where 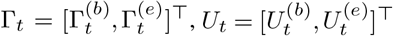 and the superscripts distinguish driver (*e* for event) and modulator (*b* for burst) *channels*. We define the 2 × *T* array of *T* samples of the input process 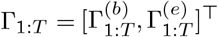 and the analogous array, *U*_1:*T*_, for the output process.

Using the definition of mutual information, and the independence of the input processes, we have

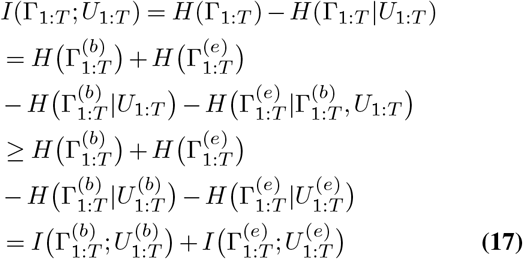

where we used the fact that entropy decreases (or is unchanged) by conditioning on another random variable (54) to get the inequality.

This shows that the mutual information in a bivariate system with independent inputs is greater than the sum of the element-wise mutual informations. It remains to check that this extends to information rates.

We define the information rate between 𝒢 and 𝒰 as the difference of entropy rates, as in (55–57):

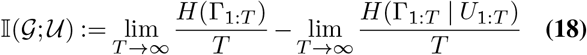

Assuming both limits exist we thus have

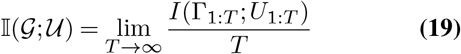

By equation 17 we have

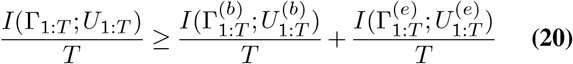

for all *T*. If the limits, with respect to *T*, of all three terms in equation 20 exist, taking the limit of both sides extends the result to information rates as desired.

### Metrics of Spike Train Irregularity

CV2 was first suggested in Ref. (28) and is defined as follows:

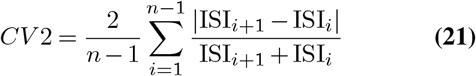

LV was developed in Ref. (29) and is given by

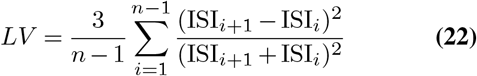

In both equations we define 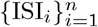 as a sequence of ISIs calculated from a single spiketrain, so that ISI_*i*+1_ follows directly after ISI_*i*_.

### Recorded data

The dataset used in Figure 1 consists of 1266 spiketrains recorded from multiple regions of the mouse brain using neuropixel probes, and has been previously published (1). We used Hartigan’s dip test (*p* ≤ 0.05) (58) to separate unimodal from multi-modal ISI distributions.

## Author contributions

EW designed the simulations, analyzed the data and conducted the theoretical analysis. AP and RN supervised the project. RN and AG designed the study.

## Acknowledgements

We thank Sam Gale, Corbett Bennett and Shawn Olsen as well as the Allen Institute for Brain Science for data sharing. RN was supported by NSERC Discovery Grant No. 06972 and CIHR Project Grant No. RN38364. EW was supported by M.Sc. scholarships from NSERC and OGS. We thank Maia Fraser for helpful discussions.

## Code Availability

https://github.com/nauralcodinglab/zeke_msc

## Supplementary Note A: Correlation theory

Correlation theory (52) is a recently proposed method to estimate the information rate of neural spiketrains. We employed correlation theory as a second method to evaluate burst coding in neural populations but, in doing so, found that this method is only a lower bound on the total information.

To prove that correlation theory calculates the full information rate in spike trains, Dettner et al. (2016) (52) used the result, due to David Brillinger, that if a time series, *Y*_*t*_, is mean-zero, finite memory and stationary, it has Fourier components that are asymptotically (59) distributed as follows

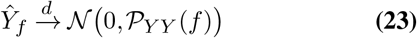

where 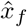 denotes the discrete Fourier transform of time series *x*_*t*_ at frequency *f, 𝒫* _*Y Y*_ the 2 x 2 diagonal matrix whose non-zero elements are the power spectrum of *Y*, and *𝒩* denotes the bivariate normal distribution. Critically, it is further assumed that if one can show empirically that the Fourier transforms of the output time series given the input time series, *Y x,t* where *X*_*t*_ denotes input time series, are asymptotically distributed with some mean and variance then the information rate is given by the correlation theory equation

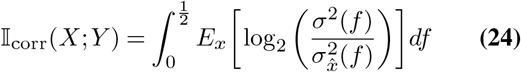

where the expected value is over sample paths of *X, σ*^2^(*f*) and 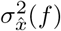 are the variances of the real and imaginary parts of *Ŷ* and variance of the real and imaginary parts of 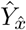, respectively. Note that one must compare the mean-subtracted time series because otherwise the moments of the Fourier transforms at *f* = 0 will not be defined.

If one can show that there exist time series *X* and *Y* where *Ŷ* _*x*_ is asymptotically normal such that the above expression fails to calculate the information rate of *X* and *Y* then correlation theory is incorrect for stationary, finite memory time series in general. We will show that such time series exist.

Let *X*_*t*_ ∼ 𝒩 (0, 1), *𝒩* _*t*_ ∼ *χ*^2^(1) such that *N*_*t*_ and 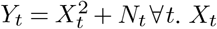 and *Y*_*t*_ are thus trivially jointly stationary and finite memory. Now assume that each of 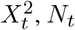 and *Y*_*t*_ are shifted so that their means are zero, allowing for Fourier transforms that are defined even at the zero frequency. Their information rate is given thus:

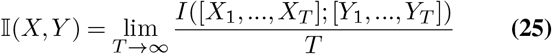

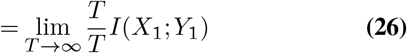

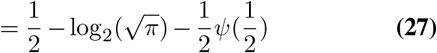

where we have used the expression for the entropy of a chi squared distribution in units of bits and the fact that shifting the location of a distribution does not effect its variance or entropy.

Now we will compute the value of the correlation theory expression. By Brillinger’s theorem the Fourier transforms of the output time series are described by equation 23. This means that *σ*^2^(*f*) = *𝒫*_*Y*_ (*f*). This is the power spectrum of a stationary time series so it is equal to the Fourier transform of the autocorrelation of *Y*_*t*_. Because *Y*_*t*_ are iid samples the autocorrelation is zero everywhere except at *τ* = 0, where it is given by the correlation function for a chi square random variable, *Z*, with 2 degrees of freedom; thus *𝒫*_*Y*_ (*f*) = *E*(*Z*^2^) = 6 ∀*f*.

By linearity of the Fourier transform, the variance of the transform of *Y*_|*x,t*_ (is

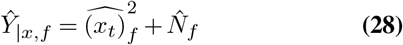

thus the variance of the real and imaginary parts of *Ŷ*_|*x,t*_ are fully described by those of 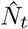. This means that *Ŷ*_|*x,t*_is asymptotically normal by the asymptotic normality of the noise process. Also by Brillinger’s proof these are equal to the power spectrum of the noise process. Because the noise process is stationary this is simply the Fourier transform of the autocorrelation function, which is zero everywhere except for zero time lag, by iid sampling, where it is equal to the correlation of a chi square random variable with one degree of freedom; thus *V* (*Ŷ*_|*x,t*_) = 3 ∀ *f*. Taken together with *𝒫*_*Y*_ (*f*) and equation 24, we get

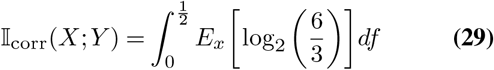

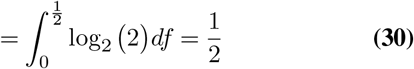

completing our counter argument. We expect that the issue is that correlation theory assumes convergence in entropy will follow from convergence in distribution, which is not always true (60).

We have shown here that correlation theory does not estimate information rate for stationary, finite memory time series in general. For specific time series, e.g. neural spike trains, the required entropy convergence may indeed occur, but, to our knowledge, this remains to be proven. Regardless, correlation theory is equal to Stein’s method in the case of a linear transfer function and is thus an estimator for information from linear decoding.

## Supplementary Note B: Supplementary figures

